# Near-Infrared Photothermal Ablation of Biofilms using Protein-Functionalized Gold Nanospheres with a Tunable Temperature Response

**DOI:** 10.1101/2023.08.12.553096

**Authors:** Dhanush L. Amarasekara, Chathuri S. Kariyawasam, Madison A. Hejny, Veeresh B. Torgall, Thomas A. Werfel, Nicholas C. Fitzkee

## Abstract

Temperature-responsive nanostructures with high antimicrobial efficacy are attractive for therapeutic applications against multi-drug-resistant bacteria. Here, we report temperature-responsive nanospheres (TRNs) that are engineered to undergo self-association and agglomeration above a tunable transition temperature (T_t_). Temperature-responsive behavior of the nanoparticles is obtained by functionalizing citrate-capped, spherical gold nanoparticles (AuNPs) with elastin-like polypeptides (ELPs). Using protein design principles, we achieve a broad range of attainable T_t_ values and photothermal conversion efficiencies (*η*). Two approaches were used to adjust this range: First, by altering the position of the cysteine residue used to attach ELP to the AuNP, we attained a T_t_ range from 34-42 °C. Then, functionalizing the AuNP with an additional small globular protein, we were able to extend this range to 34-50 °C. Under near-infrared (NIR) light exposure, all TRNs exhibited reversible agglomeration. Moreover, they showed enhanced photothermal conversion efficiency in their agglomerated state relative to the dispersed state. Despite their spherical shape, TRNs have a photothermal conversion efficiency approaching that of gold nanorods (*η* = 68±6%), yet unlike nanorods, the synthesis of TRNs requires no cytotoxic compounds. Finally, we tested TRNs for photothermal ablation of biofilms. Above T_t_, NIR irradiation of TRNs resulted in a 10,000-fold improvement in killing efficiency compared to untreated controls (p < 0.0001). Below T_t_, no enhanced anti-biofilm effect was observed. In conclusion, engineering the interactions between proteins and nanoparticles enables the tunable control of TRNs, resulting in a novel, anti-biofilm nanomaterial with low cytotoxicity.

**TOC Image:** 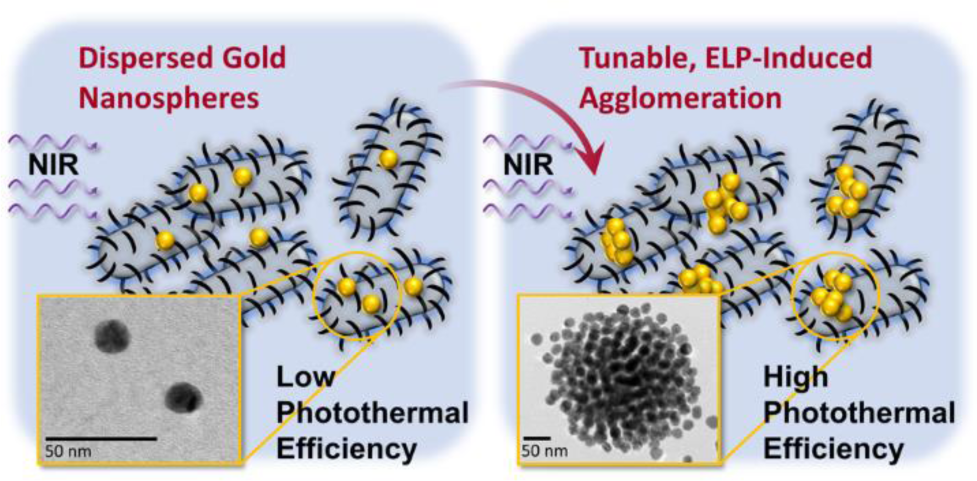

## Introduction

Gold nanoparticles (AuNPs), such as nanospheres, nanorods, nanocages, and nanostars, are readily utilized in biomedical applications due to their highly efficient photothermal conversion and biocompatibility.^1–9^ The optical properties, size, and shape of the nanoparticle ultimately determine the specific therapeutic application.^10^ Photons in the near-infrared region (NIR, 800 – 1100 nm) readily penetrate soft tissues; therefore, recent efforts have sought the ability to optimize AuNP absorption in the NIR region.^11^ For example, gold nanorods (AuNRs) have been successfully used for in vivo anticancer treatment by Dickerson *et al*. using their strong longitudinal NIR absorbance.^12^ AuNRs offer many advantages, including NIR tunability. Moreover, the photothermal conversion efficiency (η) is high, which makes treatment of cancers and other ailments possible with low concentrations of nanoparticles. For example, Qin *et al.* obtained a photothermal conversion efficiency of 65% for nanorods with an aspect ratio of 3.8 (D = 10.6 nm, L = 40 nm), meaning that 65% of absorbed photon energy could be harvested to generate heat.^13^ However, AuNRs have several drawbacks. Their synthesis requires multiple steps, and the compounds used in their manufacture can induce cytotoxicity.^14,15^ These factors reduce the suitability for AuNRs in the clinic. Compared to AuNRs, gold nanospheres (AuNSs) are less cytotoxic and can be synthesized using only chloroauric acid and citrate in a one-pot synthesis.^16^ But AuNSs are not optimal for photothermal therapeutic (PTT) applications, because their localized surface plasmonic resonance (LSPR) lies in the visible region.^1,2,17^

Self-assembled or agglomerated systems of small AuNSs can be employed to increase photothermal conversion efficiency. This is due to the plasmonic coupling effect between adjacent AuNSs (Supporting Information, **Fig. S1**), which increases the extinction in the NIR.^2,3,18^ Most of these studies have used biomacromolecules to functionalize the gold surface, improving biocompatibility and optimizing PTT responsiveness. Elastin-like polypeptides (ELPs) are an attractive option for this purpose. ELPs represent a class of synthetic, five-residue peptide repeats based on human elastin.^19–21^ These repeats typically have the form VPGXG, where X represents a guest residue.^22^. ELPs undergo protein phase separation at elevated temperatures,^23–25^ allowing them to drives self-association of nanoparticles above a tunable transition temperature (T_t_). The transition temperature is tunable and depends on the identity and overall composition of the guest residue.^26–28^

Nath *et al*., Higashi *et al*., and Sun *et al*. have previously used ELPs with this strategy to prepare temperature-responsive gold nanoparticles (AuNP@ELP).^2,17,29^ These three approaches differ slightly, but they suffer from some common limitations. The earliest approach was developed by Nath *et al.,* who demonstrated that AuNP@ELP could self-associate reversibly and developed the initial characterization of these materials. Their ELP construct consisted of 180 pentapeptide repeats, and AuNP functionalization was performed by non-covalent adsorption of ELP on to a self-assembled monolayer of mercaptoundecanoic acid.^29^ Higashi *et al.* synthesized a four-repeat ELP construct using solid-phase synthesis, and they used the dithiolane ring of an attached lipoic acid to covalently functionalize AuNPs. With only four repeats, the T_t_ was fairly high, 40 °C, and the authors did not explore approaches to modulate the T_t_.^17^ Most recently, Sun *et al*. explored the photothermal therapy potential of ELP-functionalized AuNPs. A 26 kDa ELP was used, and an N-terminal cysteine residue allowed attachment to AuNPs.^2^ The construct showed promise for photothermal ablation of cancers in a mouse model, but the conversion efficiency η was only marginally higher than PEG-AuNPs, 30% vs 21%. They were able to tune the temperature response of the AuNP@ELP by changing the surface density of ELP on the AuNP surface,^2^ but the observed temperature range was narrow (about 2℃) and the T_t_ was estimated to be 22 ℃, which is not optimal for intravenous injection of these nanoparticles.^30,31^ ELP-functionalized AuNPs show a great deal of promise, but challenges remain with respect to tunability of T_t_ and the optimization of η.

An obvious potential application of highly tunable AuNP@ELP is treatment of biofilms on medical devices. In recent years, medical device-related infections have become a serious problem, and these infections are a leading cause of death around the world.^32–34^ Pathogenic microorganisms like *S. epidermidis* are known to cause persistent infections by forming biofilms.^33,35–38^ Bacteria in these biofilms are protected from antibiotic penetration by a layer of extracellular polymeric substances (EPS), making infections more challenging to treat.^36,39^ Generally speaking, PTT-enabled nanomaterials are ideal for treating biofilms because heat can readily penetrate EPS when antibiotics cannot.^39–41^ Heat can destroy the structure of biofilm by irreversible cell damage, through disrupting membranes permeability, denaturing proteins/enzymes, and inducing bacterial death.^42,43^ However, values of η for existing functionalized materials used for biofilm treatment range from 20 to 30%.^2,3,32^ Higher photothermal conversion efficiencies and more tunable nanomaterial behavior would enable more flexibility in the delivered nanoparticle dose, applied NIR energy, and the range of tissues that could be treated. An ELP-functionalized platform based on spherical AuNPs would be ideal for this application, provided sufficient tunability were attainable. Here, we describe a facile approach for the preparation of, tunable, thermally triggered AuNP@ELP particles novel intelligent temperature-responsive nanospheres (TRNs) for antibacterial applications. In this study, two different thermally sensitive elastin-like polypeptides (ELP) were prepared and site-specifically conjugated to AuNPs to form TRNs (**Fig. 1A**). While these AuNPs exhibit photothermal conversion over a broad range of temperatures, η is significantly elevated above the T_t_, which can be programmed over a broad temperature range between 34 – 49 °C. This flexibility is enabled by controlling the ELP densities on the AuNP surface and by modulating the ELP attachment site (**Fig. 1B**). Our approach allows us to approach values of η up to 68%, rivaling the values observed for AuNRs. Under NIR irradiation above the T_t_, the self-associated TRNs within *S. epidermidis* (gram-positive) and *E. coli* (gram-negative) biofilms exhibit outstanding photothermal ablation properties, whereas the dispersed TRNs below the T_t_ show minimal damage (**Fig. 1C**). These TRNs are a promising new tool for the selective photothermal treatment of infections and demonstrate that spherical nanoparticles can be highly effective for photothermal conversion.

**Figure 1.**
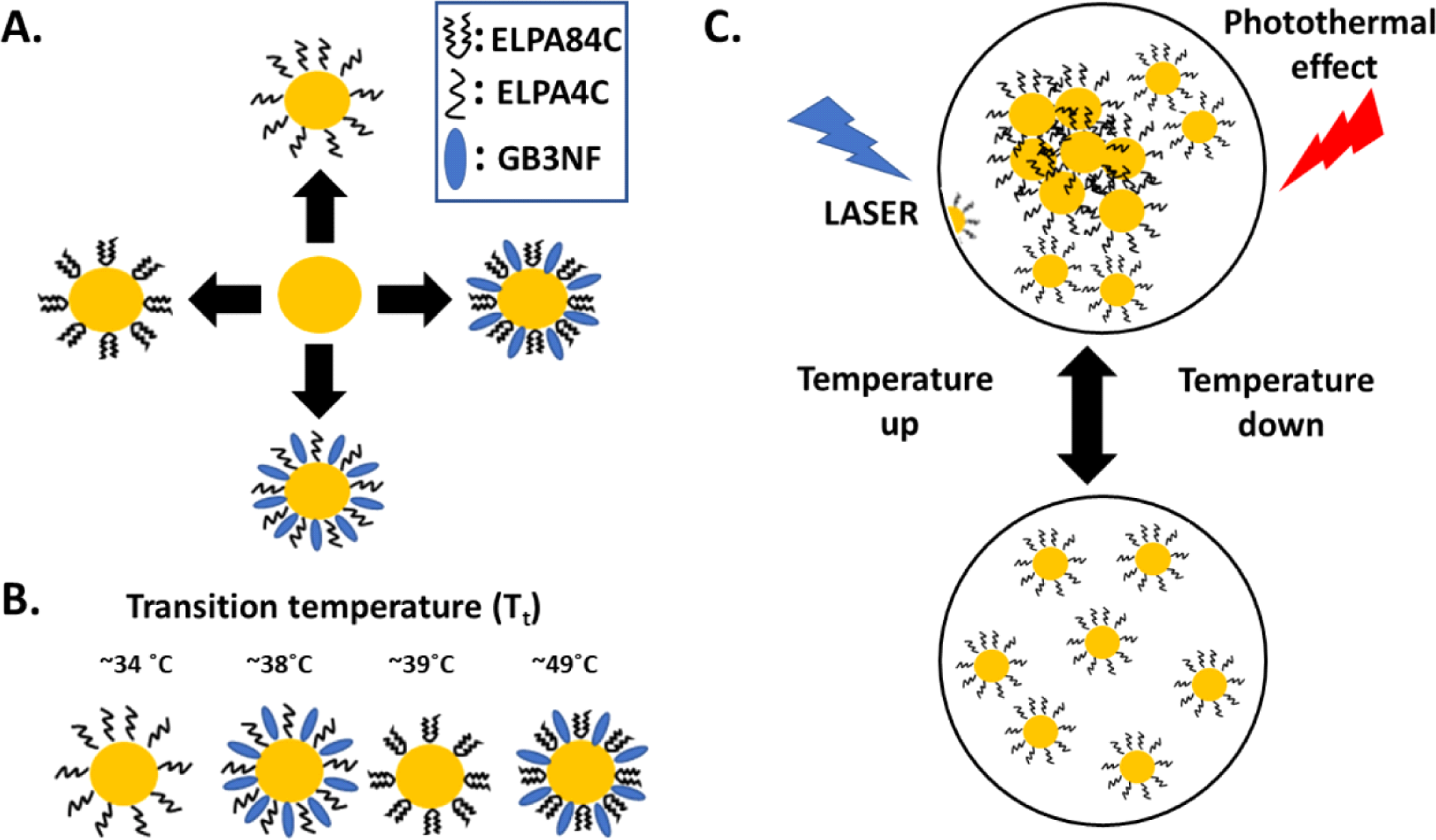
Functionalizing AuNPs with Elastin-Like Polypeptides. (A) Schematic diagram of surface modification process of AuNPs with ELP and GB3NF. (B) Change in transition temperature with surface modification. (C) Temperature responsive behavior of TRNs.

## Results and Discussion

### Towards Temperature-Responsive Nanospheres: Synthesis and Characterization of AuNP@ELP

As illustrated in **Fig. 1A**, we utilized elastin-like peptides (ELPs) to functionalize gold nanoparticle (AuNP) surfaces, creating temperature-responsive nanospheres (TRNs). While gold nanorods have excellent photothermal conversion efficiency in the NIR region,^13,44,45^ we hypothesized that ELP-induced agglomeration would induce a similar enhancement in photothermal properties when AuNP@ELP nanospheres were irradiated with NIR light. For the synthesis, 15 nm diameter spherical AuNPs were used (Supporting Information, **Figs. S2-4**). Two single-site cystine variants were engineered, one with a cysteine residue at the N-terminus (ELPA4C) and another with a central cysteine residue (ELPA84C). These cysteines were used to conjugate ELP to AuNPs through a gold-thiol linkage (Supporting Information, **Fig. S5**). Most of the guest residues in the ELP were valine, chosen to lower the transition temperature (T_t_) (Supporting Information, **Fig. S5**).^26^ Both ELP constructs were able to reproduce temperature-induced liquid-liquid phase separation (LLPS) (Supporting Information, **Fig. S6**), and the T_t_ was concentration-dependent (Supporting Information, **Fig. S6,** inset).^35^ We explored two molar ratios of AuNP to ELP, a low ratio (1:200) and a high ratio (1:1000). Interestingly, while AuNP@ELP exhibits a larger hydrodynamic diameter (DH) than non-functionalized AuNPs, the molar ratio did not dramatically affect the observed particle size. For AuNP@ELPA4C, DH increased from 17 nm for non-functionalized AuNP to 43 nm for 1:200 AuNP@ELPA4C, but higher ELP ratios did not produce significantly larger nanoparticles (**Fig. 2A**, orange vs. red curves; **Table 1**). A similar trend was observed for AuNP@ELPA84C, where DH was not observed to increase as a function of molar ratio (**Fig. 2B**, **Table 1**). The DH values of the AuNP@ELPA4C were higher than those of AuNP@ELPA84C. This may reflect the different tethering mode of ELPA4C vs. ELPA84C: Because ELPA4C is bound near the N-terminus, it is always relatively extended. On the other hand, ELPA84C is constrained by its centralized tethering site. UV-Vis extinction spectra also confirmed the successful functionalization of ELP on the AuNP surface and followed a similar trend (Supporting Information, **Fig. S7**).

**Figure 2.**
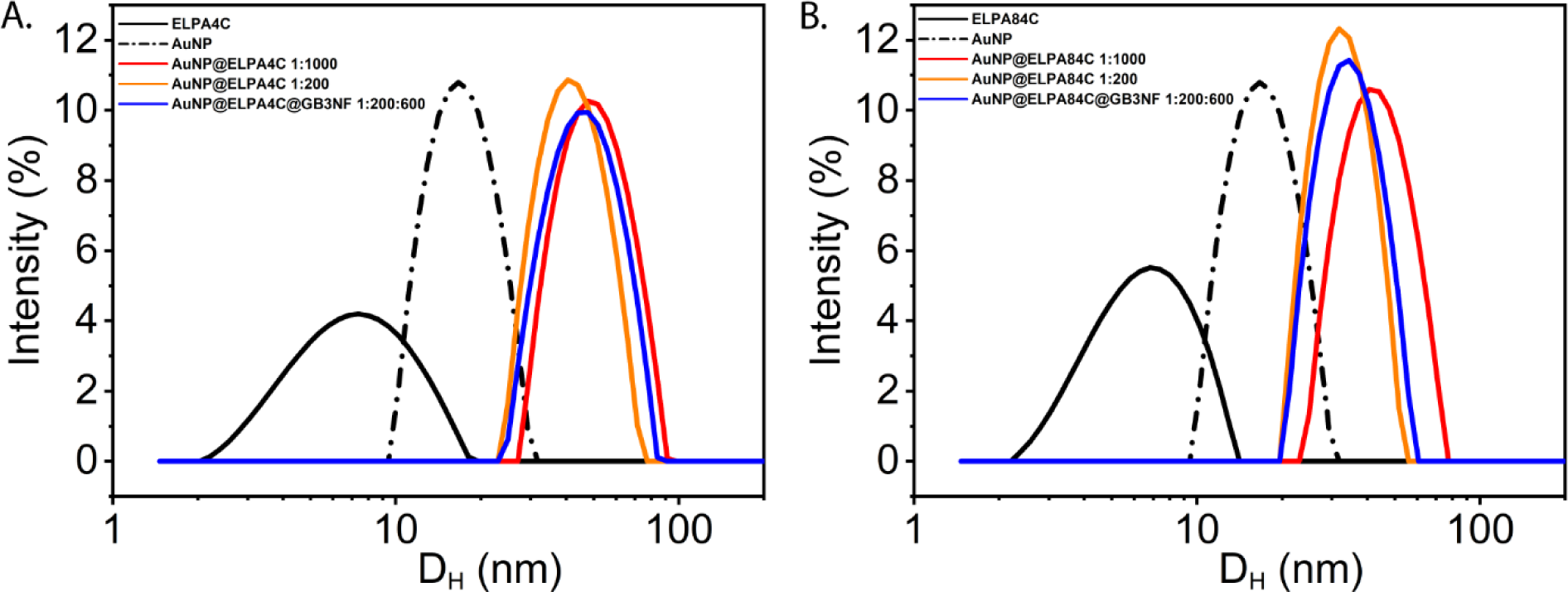
DLS profiles of ELP, AuNP, and TRNs. DH denotes hydrodynamic diameter. (A) ELPA4C functionalized TRNs. (B) ELPA84C functionalized TRNs.

**Table 1.**
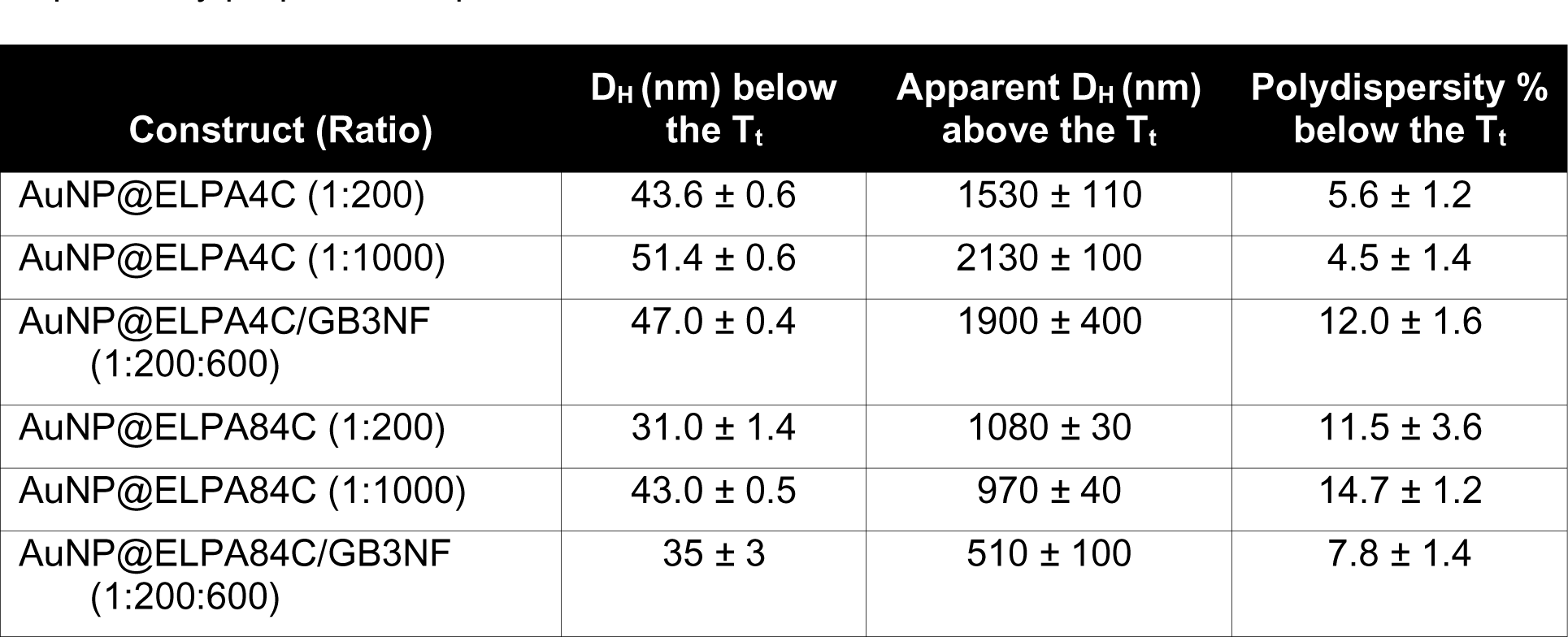
Summary of hydrodynamic diameter (DH) for TRNs below and above their transition temperatures (T_t_). Errors represent the standard error of the mean from at least three independently prepared samples.

| Construct (Ratio) | $D_H$ (nm) below the $T_t$ | Apparent $D_H$ (nm) above the $T_t$ | Polydispersity % below the $T_t$ |
| --- | --- | --- | --- |
| AuNP@ELPA4C (1:200) | $43.6 \pm 0.6$ | $1530 \pm 110$ | $5.6 \pm 1.2$ |
| AuNP@ELPA4C (1:1000) | $51.4 \pm 0.6$ | $2130 \pm 100$ | $4.5 \pm 1.4$ |
| AuNP@ELPA4C/GB3NF (1:200:600) | $47.0 \pm 0.4$ | $1900 \pm 400$ | $12.0 \pm 1.6$ |
| AuNP@ELPA84C (1:200) | $31.0 \pm 1.4$ | $1080 \pm 30$ | $11.5 \pm 3.6$ |
| AuNP@ELPA84C (1:1000) | $43.0 \pm 0.5$ | $970 \pm 40$ | $14.7 \pm 1.2$ |
| AuNP@ELPA84C/GB3NF (1:200:600) | $35 \pm 3$ | $510 \pm 100$ | $7.8 \pm 1.4$ |

In addition to changing the tethering location, we also changed the surface density of ELP on the AuNP surface. Our approach was to add an inert, globular protein to dilute the number of ELP molecules bound to AuNPs. We used a cysteine-containing variant of the third IgG-binding domain from streptococcal protein G (GB3NF, Supporting Information, **Fig. S5**).^46^ This small GB3NF protein is a 56-residue GB3 domain modified to contain a Cys-residue and a “null fusion” containing a disordered C-terminal linker. The ratios of AuNP, ELP, and GB3NF were varied during the AuNP surface modification reaction. We varied these from 1:200:200 to 1:200:600. The AuNP@ELPA4C@GB3NF (1:200:600) nanoparticles had an DH of 47 nm, and the AuNP@ELPA84C@GB3NF (1:200:600) nanoparticles had a DH of 35 nm (**Fig. 2** and **Table 1**)

### TRNs reversibly agglomerate at a tunable transition temperature (T_t_)

Next, we sought to examine the properties of AuNP@ELP at different temperatures. Successful synthesis should result in TRNs that agglomerate above T_t_ and exhibit colloidal stability below T_t_. Moreover, the T_t_ should be tunable depending on the type of ELP and number density on the AuNP surface. Temperature responsiveness of the synthesized TRNs was confirmed by measuring extinction at 600 nm (turbidity), and DH using DLS, both as a function of temperature (**Fig. 3A**). The DLS experiments revealed a significant increase in apparent hydrodynamic diameter with increasing temperature (Summarized in **Table 1**), and this diameter returned to < 100 nm after the temperature was lowered. The plasmonic absorption peak has shifted to a longer wavelength implying that the particles agglomerated to larger clusters when the temperature was increased. The temperature at which a sharp increase was observed was taken as the T_t_. Below the T_t_, the TRNs in suspension have an average interparticle distance greater than the colloidal diameter. Upon raising the temperature, the adsorbed ELP on the surface undergoes self-association, resulting in large, reversible nanoparticle agglomerates (Supporting Information, **Fig. S8**).^29^

**Figure 3.**
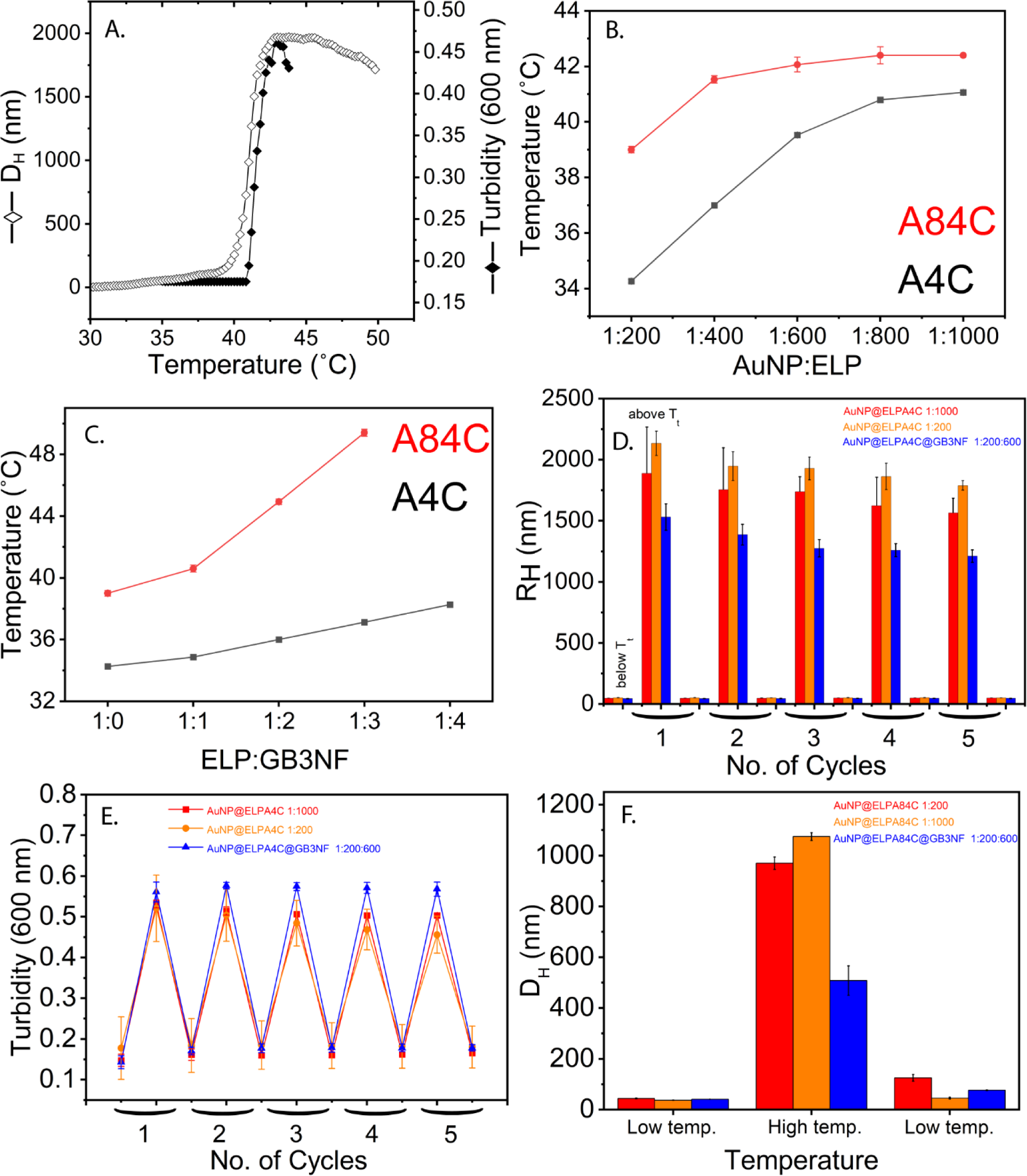
Characterizing temperature response of AuNP@ELP nanoparticles. (A) Examination of ELP agglomeration using DLS and turbidity. (B) The ELP concentration dependence of the phase transition temperature (T_t_) of TRNs with A84C (red) and A4C (black). (C) T_t_ of TRNs as a function of ELP:GB3NF ratio for A84C (red) and A4C (black). (D) DLS measurements on ELPA4C-functionalized TRNs (all at 20 nM particles, or 240 ppm Au) for 5 cycles. A reversible, temperature dependent agglomeration is observed. (E) UV-Vis turbidity measurements corresponding to (D). (F) DLS measurements on ELPA84C-functionalized TRNs for one cycle (20 nM particles, or 240 ppm Au). Only a single cycle was used because it took significantly longer for these TRNs to regain colloidal stability (∼24 hr). Error bars represent the standard error of the mean for at least three independently prepared samples.

**Fig. 3B** demonstrates that T_t_ values can be raised by increasing the stoichiometry of ELP on the surface of AuNPs. For both A4C and A84C, a tunable T_t_ was observed, with T_t_ increasing as the ELP ratio was increased. Previously, it was observed that increasing the PEG stoichiometry on AuNPs reduced its flexibility.^47^ Here, a similar effect likely reduces the flexibility of ELP, whereby increased packing could sterically occlude interactions that may lead to intermolecular association, such as turn-turn interactions.^23,24^ This occlusion would increase the temperature at which association occurs, as we observe (**Fig. 3B**). With ELP alone, T_t_ can be tuned over a range of 34-42 °C. The range of T_t_ values can also be adjusted by adding GB3NF to the surface as described above. Starting with our lowest AuNP:ELP ratio (1 AuNP per 200 ELPs), we added an increasing amount of GB3NF to further lower the amount of ELP on the surface. This reduces the surface density of ELP, making intermolecular association more difficult, which increases the T_t_. Thus, we observed that T_t_ can be increased at high ELP densities because of steric occlusion, but it can also be increased at low ELP densities because of a dilution of ELP. Dilution of ELP is known to increase the T_t_,^28^ and it is reasonable that too little ELP on the surface will make self-association of AuNPs more difficult. By changing both the ratio of ELP and GB3NF we were able to obtain a final range of 34-49 °C for T_t_. Interestingly, dilution with GB3NF on the surface appears to be more effective at spanning a broader range of temperatures, although using ELP alone may be beneficial for more precise tuning of T_t_ over a narrower range.

The reversibility of the thermally induced assembly and disassembly was also confirmed by turbidity and size measurements (**Figs. 3D-F**). Based on our design, AuNP@ELP should agglomerate above T_t_ and regain colloidal stability when the temperature is lowered. When the solution temperature was returned to room temperature, the turbidity returned to its original value, implying that the clusters were disassembled. This behavior was also seen in size measurements by DLS. This reversible assembly and disassembly behavior remained reproducible over five repeated temperature cycles for all AuNP@ELPA4C constructs (**Figs. 3D-E**). However, the A84C constructs took much longer to return to solution after agglomeration (approximately 24 h, **Fig. 3F**). This increased time reflects strong interactions between AuNP@ELPA84C particles, although the physical mechanism for the longer cycling time remains unclear.

### TRNs can exhibit a comparable photothermal conversion efficiency to gold nanorods

Next, we sought to exploit the plasmonic peak shift induced by TRN agglomeration for potential applications in photothermal therapy.^33^ AuNP self-association has been used to change the photothermal absorption efficiency;^2,32^ the TRN we developed enable a highly controllable system whereby photothermal heating could be used to kill bacteria or ablate unwanted cancer cells.^4,6,33,41^ At elevated temperatures or in agglomerated states, TRNs should have a higher photothermal effect than at lower temperatures or in non-agglomerated states. To demonstrate this, we designed an experimental setup that could simultaneously monitor thermal changes using a camera upon NIR laser irradiation (**Fig. 4A**). At first, we studied the photothermal effect of each TRN at different gold concentrations, keeping the laser power constant (1.6 W cm^-2^) and starting at the ELP T_t_ (**Fig. 4B** and Supporting Information, **Fig. S9**). For 50 nM particle concentration (600 ppm total Au) AuNP@ELPA84C, with a 1:200 ratio of AuNP to ELP and a starting T_t_ of 39 °C, the solution temperature increased by 29 °C after five minutes of 1.6 W cm^-2^ irradiation (**Fig. 4B**, orange bars). In contrast, AuNPs functionalized with an inert 5,000 molecular weight thiolated polyethylene glycol (5K PEG-SH, or AuNP@PEG5K) at 35 °C showed a smaller effect, increasing only 16 °C (Supporting Information, **Fig. S10**). Even at a particle concentration of 20 nM (240 ppm Au), all TRNs showed a marked increase in solution temperature with increasing laser power (**Fig. 4C** and Supporting Information, **Fig. S11**). Furthermore, the temperature increase became smaller when the TRNs were below T_t_ in a non-agglomerated state. In those solutions, the temperature increase reached only about 7-12 °C after 5 minutes of exposure, resulting in a large difference in temperature change (ΔΔT) between the agglomerated and non-agglomerated states (**Fig. 4D** and Supporting Information, **Fig. S12**). More specifically, a solution containing AuNP@ELPA84C 1:200 showed a ΔΔ*T* of 10.7 ± 1.1 °C in its agglomerated state relative to its non-agglomerated state (**Fig. 4D**). The photothermal conversion efficiency (ƞ) of AuNP@ELPA84C 1:200 was calculated to be around 68±6% (**Fig. 4E** and Supporting Information, Methods), which was higher than that of AuNP@PEG5K (31±5%, Supporting Information Methods).

**Figure 4.**
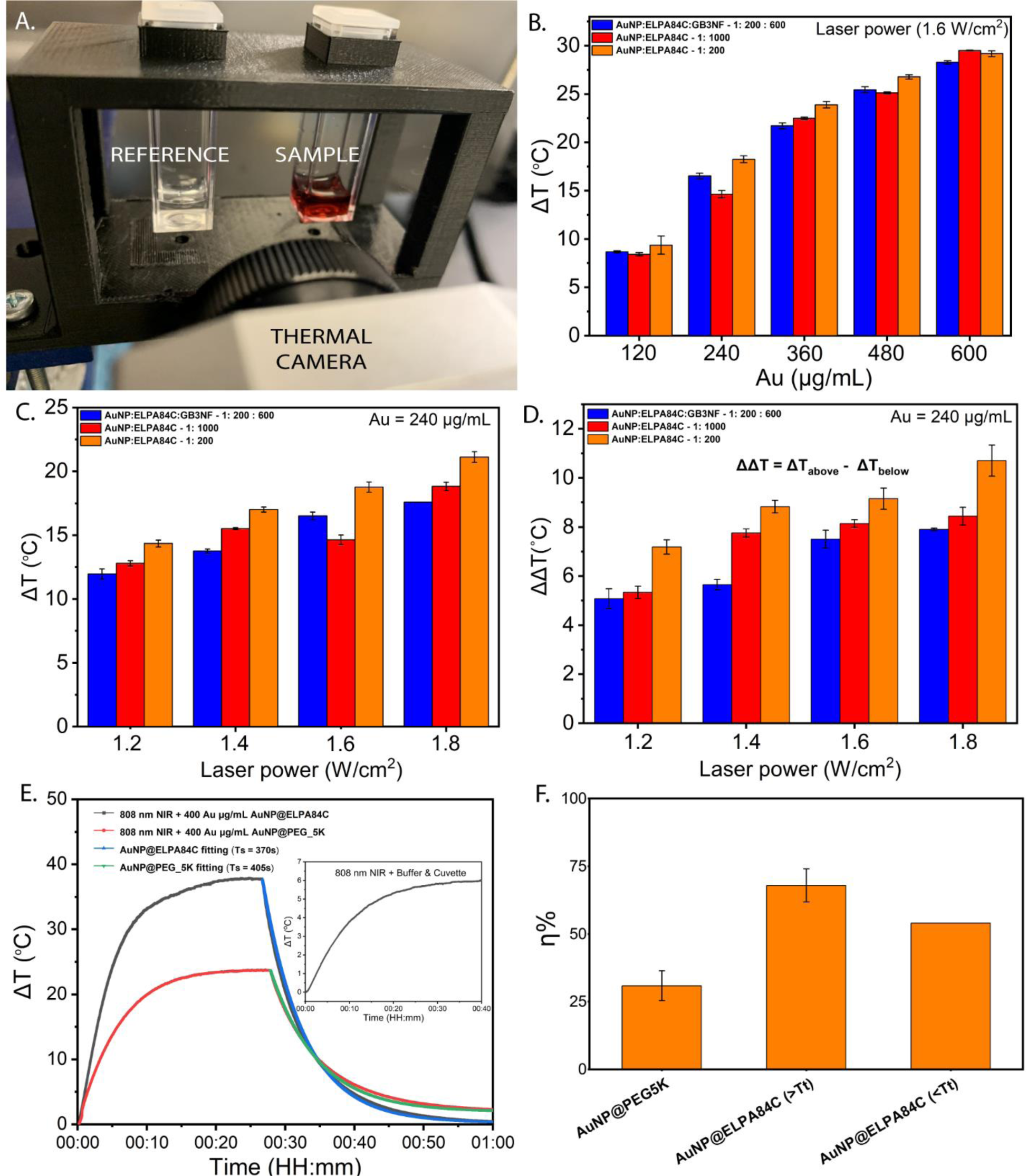
(A) Digital image of the experimental setup used for the measurements of photothermal effect. (B) The gold concentration dependence and (C) laser power dependence on TRNs solution temperature change when exposed to laser irradiation 808 nm (1.6 W cm^-2^, 5 min), after incubation for 15 min at their transition temperatures (T_t_). (D) The difference in solution temperature change (ΔΔ*T*) at their agglomerated state vs. the non-agglomerated state. (E). Photothermal effect of the irradiation of the 20 nM (240 ppm total Au) aqueous solution of AuNP@ELPA84C 200:1 and AuNP@PEG5K with a 1.8 W cm^-2^, 808 nm laser. 20 mM HEPES + 100 mM NaCl with a 1.8 W cm^-2^, 808 nm laser (inset). (F.) Change in photothermal conversion efficiency (ƞ) above and below T_t_ of AuNP@ELPA84C 1:200. The value of η below T_t_ is estimated from the biphasic temperature data starting at 22 °C (Supporting Information, **Fig. S13**). Error bars represent the standard error of the mean for at least three independently prepared TRN samples.

One complication in measuring ƞ in these materials is the changing nature of the solution as nanoparticle agglomeration evolves. Within the first five minutes of irradiation, ELP-coated AuNPs are in the earliest stages of agglomeration, and most particles remain suspended in solution. As time passes, more agglomeration occurs, leading to further increase in Δ*T* and η. This is demonstrated by a biexponential increase in temperature when AuNP@ELPA84C (1:200) is irradiated well below its T_t_ at 22 °C (Supporting Information, **Fig. S13**). An increase occurs that results from initial irradiation, followed by a second increase once agglomeration begins to accelerate. The estimated ƞ for these two regimes is 54% (below T_t_) vs. 69% (above T_t_), and the average value is somewhere in between (**Fig. 4F** and Supporting Information Methods). Presumably, η would decrease again as nanoparticle agglomerates began to sediment, lowering the concentration in solution. Thus, the actual η is time dependent, but even in the range bracketed above, the AuNP@ELPA84C 200:1 construct has a photothermal efficiency in the NIR comparable to some gold nanorods (65%),^13^ which is surprising given its spherical shape. All of these data suggest that TRNs synthesized in this study are highly efficient photothermal transducers and show comparable photothermal effect to gold nanorods in a similar concentration range.^6^

### TRNs are biocompatible and possess tunable in vitro antibiofilm activity when irradiated by NIR light

Next, we investigated whether the AuNP@ELP constructs were active against bacterial biofilms and biocompatible with human cell culture. To evaluate the bactericidal effect, TRN solutions were applied to two different biofilms formed under static conditions from *E. coli* (gram negative, strain ATCC 25992) and *S. epidermidis* (gram positive, strain 1301).^37^ Here, we focused on the TRN with the highest thermal response, AuNP@ELPA84C in an AuNP:ELP ratio of 1:200. Biofilms were grown for 48 hr at 37 °C in cell culture plates, after which excess media was removed and the biofilm was washed with PBS. After this, the biofilm was treated with 20 nM particle concentration of TRN (240 ppm total Au, or 96 μg Au per well), then immediately irradiated by NIR light (**Fig. 5**). We then determined the number of viable colonies by collecting the biofilm and plating serial dilutions (CFU mL^-1^). This simple test determines whether the thermal response of our engineered nanoparticles is sufficient to kill bacterial cells embedded in biofilms, where EPS and other biomass can protect bacteria from traditional antibiotic exposure. We sought to use a low concentration to minimize the AuNPs needed. The concentration of TRNs used here is approximately twice what was used by Hu *et al*.,^32^ who used 110 ppm total Au in their work. Hu *et al.* found that pH-responsive AuNPs raised the temperature of a MRSA biofilm by 33 ℃ in 10 min under 0.94 W cm^-2^ NIR laser. The two experiments are not directly comparable because of differing applications and functionalization, but they demonstrate that TRNs can obtain similar performance to other platforms using functionalized AuNPs.

**Figure 5.**
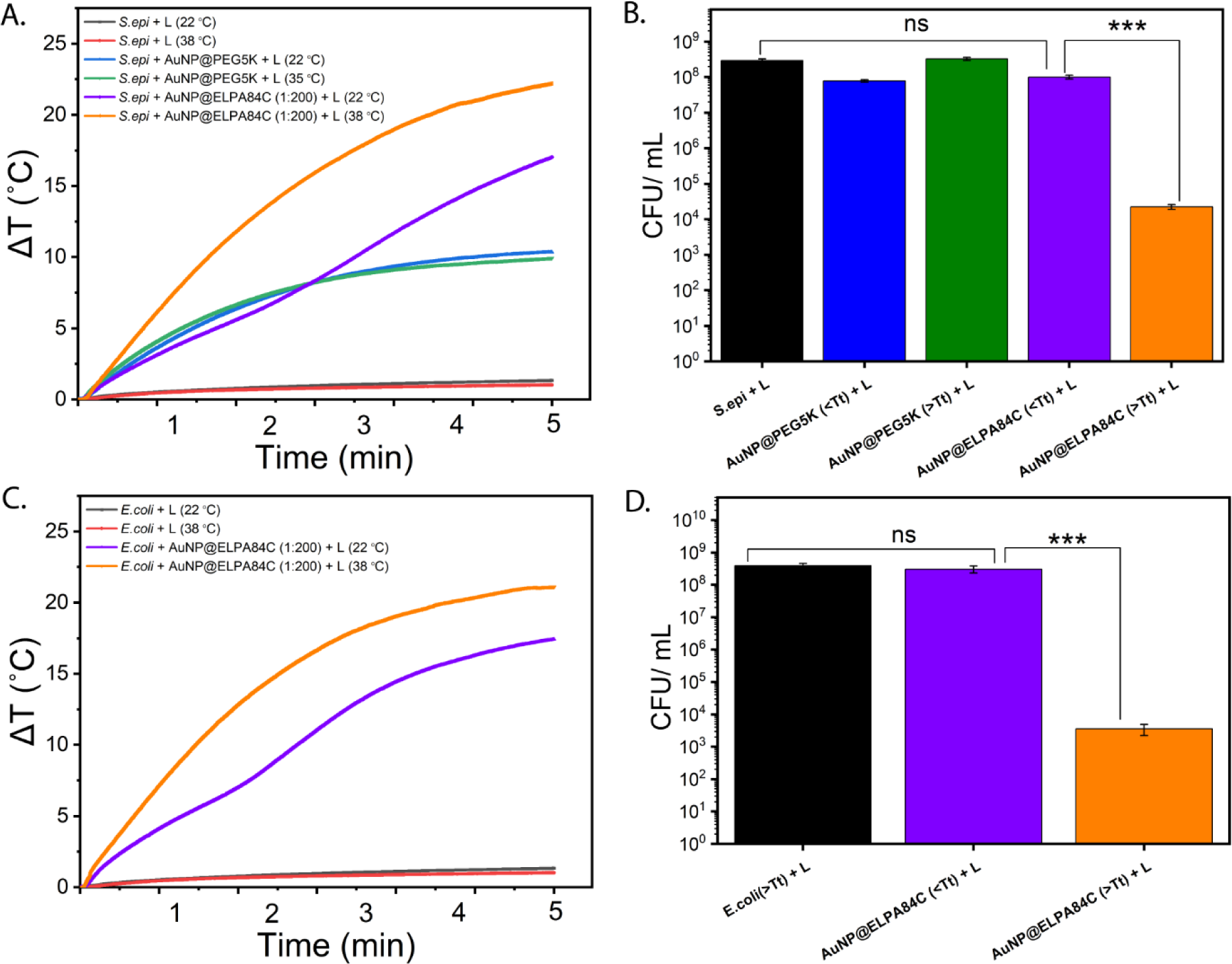
(A). Temperature evolution curves of *S. epidermidis* biofilm, AuNP@PEG5K in *S. epidermidis* biofilm, AuNP@ELPA84C (1:200) solutions in *S. epidermidis* biofilm under NIR irradiation at below and above transition temperature. (B). Number of *S. epidermidis* bacterial colonies at below and above transition temperature, after a negative control, AuNP@PEG5K, and AuNP@ELPA84C (1:200) with and without NIR irradiation for 5 min. (C). Temperature evolution curves of *E. coli* biofilm, AuNP@ELPA84C (1:200) solution in *E.coli* biofilm under NIR irradiation at below and above transition temperature. (D). Number of *E. coli* bacterial colonies at below and above transition temperature, after treated with nothing, and AuNP@ELPA84C (1:200) under NIR irradiation for 5 min. Significance was assessed using one-way ANOVA with Tukey’s multiple comparisons test (n.s., not significant; ***, p < 0.0001). Error bars represent the SEM of three experiments.

We confirmed the presence of *E. coli* and *S. epidermidis* biofilms on treated polystyrene well plates using a crystal violet stain and scanning electron microscopy (Supporting Information, **Fig. S14**). Once irradiated by NIR light, the temperature of the *S. epidermidis* and *E. coli* biofilm containing AuNP@ELPA84C increased rapidly and showed a maximum temperature of approximately 60 °C (**Fig. 5A**). This experiment, performed at the T_t_ (39 °C) for AuNP@ELPA84C shows the excellent photothermal effect of TRNs even in the presence of biofilm. Many bacterial proteins become denatured above 50 °C, leading to cell death.^2^ The number of viable bacteria decrease significantly after NIR treatment relative to controls with no nanoparticles, non-irradiated PEGylated AuNPs, or with NIR-irradiated PEGylated AuNPs (**Fig. 5B**). Bacterial viability was monitored as log10 (CFU mL^-1^) for each of the different conditions (**Fig. 5C**). Without treatment, both *E. coli* and *S. epidermidis* reached CFU mL^-1^ values of ∼10^8^ or more, reflecting a dense, healthy biofilm. After treatment with TRNs, a four-log unit reduction (99.98% reduction in cell viability) was observed in the CFU mL^-1^ compared to the controls. This surpasses the study by Hu *et al*., where an 80% reduction of MRSA biofilms was observed.^32^

Interestingly, when the temperature was lowered to 22 from 39 °C, there was no obvious change in the cell viability for AuNP@ELPA84C, reflecting the fact that ELP behaves differently above vs. below its T_t_. This observation holds for the other TRNs synthesized in this study (Supporting Information, **Fig. S15**). As noted above, we see a similar reduction in four log units when TRNs are above T_t_ in an agglomerated state; however, no significant change is observed when the temperature is below T_t_. This likely results from three factors: First, lowering the ambient temperature makes it easier for bacterial cells to survive small increases in temperature induced by photothermal heating of non-agglomerated spheres. Second, and more importantly, this behavior also reflects the reduced photothermal conversion efficiency in nanoparticle states lacking agglomerated TRNs. Very little temperature change overall is observed below T_t_, reflecting the smaller *η* for these particles (**Fig. 4F**). Spherical nanoparticles have intrinsically low photothermal conversion efficiency at the NIR frequencies used here, and this suggests that below the T_t_, TRNs behave more like traditional 15 nm nanospheres. Third, there is likely a cooperative effect above the T_t_: as TRNs begin to agglomerate, the temperature rises under continuous irradiation, triggering more agglomeration. Thus, the ELP-functionalized nanoparticles presented here exhibit a tunable anti-biofilm response: depending on the surface functionalization, TRNs can be synthesized with a variable photothermal efficiency depending on the temperature, with a tunable temperature cutoff over a range of 34-49 °C (**Fig. 3B-C**). Since higher concentrations can produce a lower T_t_,^28^ this range can likely be extended depending on the local AuNP concentration. The tunable behavior in T_t_ represents a unique feature to this class of materials.

Finally, we examined whether the TRNs altered cell viability for human embryonic kidney cells (HEK-293) in the absence of NIR irradiation. Particle concentrations of 10, 20, and 40 nM were tested (120, 240, 480 ppm total Au), covering the same range used to disrupt biofilms (Supporting Information, **Fig. S16**). As anticipated, bare citrate-capped AuNPs and AuNP@PEG5K showed no change in the viability of HEK-293 cells after incubation for 24 h with TRN. The behavior of AuNP@ELPA84C and AuNP@GB3NF were identical, confirming that, in the absence of NIR irradiation these particles biocompatible.

Together, these results demonstrate that the ELP-functionalized AuNPs presented here can photothermally ablate biofilms with several key advantages over other nanoparticle-based anti-biofilm materials. First, we are able to use low particle concentrations and relatively short, 5 min treatment times, comparable to current best-in-class anti-biofilm nanoparticles.^32^ The low concentration requirement also potentially enables applications with targeted nanoparticle delivery, since the local concentration of biofilm-targeted TRNs is likely to similar to the values used here (240 ppm). Second, ELP-functionalized AuNPs are synthesized from citrate-capped gold spheres, offering advantages in simplicity and biocompatibility. Compared to nanorods, the synthesis of 15 nm gold nanospheres is very straightforward, involving only citrate, a weak, biocompatible reducing agent. To the contrary, the surfactants commonly used to synthesize nanorods, such cetyltrimethylammonium bromide (CTAB), are cytotoxic,^48^ necessitating their complete removal from nanoparticle suspensions intended for biological applications. Even so, CTAB-coated nanorods can exhibit toxicity at higher concentrations unless additional surfactant coatings are used.^49^ Thus, the TRNs proposed here can be tuned for comparable photothermal efficiency as some gold nanorod systems (68% vs. ∼65%)^50^ without the use of potentially cytotoxic starting materials. Third, the AuNP@ELP materials presented here have an advantage in that they are switchable, exhibiting a lower photothermal efficiency below T_t_. Since T_t_ and correspondingly Δ*T* are dependent on the concentration (**Fig. 4B**),^27,28^ these AuNPs can be tuned to have relatively low photothermal conversion efficiency (similar to nanospheres) at low concentrations or lower temperatures. Thus, while nanorods remain the standard for photothermal therapies, ELP-based TRNs may be an attractive alternative, and the biofilm experiments demonstrate their potential in a realistic model. Work is ongoing to augment AuNP@ELPs with anti-fouling and biofilm targeting functionality.

## Conclusions

We synthesized temperature-responsive, ELP-functionalized spherical AuNPs using cystine based conjugation and investigated their photothermal properties, including their performance in a simple anti-biofilm assay. These TRNs agglomerate in response to elevated temperature, forming particle clusters with high photothermal conversion efficiency upon near infrared radiation. The transition temperature for agglomeration is tunable over a broad temperature range that includes physiological temperature. Surprisingly, despite their spherical shapes, the best ELP-based TRNs exhibit similar photothermal conversion efficiency to nanorods and do not require the use of cytotoxic chemicals in their synthesis. This temperature responsive self-association of the synthesized particles can be potentially employed for developing intelligent photothermal transducers for thermal ablation of biofilms. The mechanism of temperature responsive behavior is based protein phase separation behavior in synthetic ELPs, and tunability is attained by controlling the amino acid sequence and Cys residue placement. Like ELP phase separation, TRN agglomeration is reversible, providing an attractive mechanism for nanoparticle dispersal after treatment. *In vitro* evaluation of the anti-biofilm properties of the synthesized AuNP assemblies showed that the particle clusters have superior a photothermal therapeutic effect compared to non-temperature responsive particles. The TRNs presented here provide simple and general strategy for efficient and safe agents for photothermal therapy.

## Materials and Methods

### Preparation of 15 nm, Spherical Gold Nanoparticles (AuNPs)

Gold nanoparticles were synthesized using the citrate reduction method.^51–53^ The prepared AuNPs were concentrated at 6,600 × *g* for 45 min at 4°C. The concentrated samples were characterized via UV-Visible spectroscopy, dynamic light scattering (DLS) and transmission electron microscopy for size and conformity.

Particle Size Distribution (PSD) was determined by DLS (Supporting Information, **Fig. S2**). Measurements were obtained with an Anton Paar Litesizer 500 DLS system while controlling the temperature. Each sample was recorded at 20 °C, in triplicate, using the Kalliope software from the DLS intensity-weighted particle size distribution. Synthesized AuNPs showed a hydrodynamic diameter slightly above 15 nm because of associated solvent molecules on the AuNP surface. The polydispersity index was below 6%, indicating that the AuNPs are highly homogenous in size. This hydrodynamic diameter corresponds to a 15-nm AuNP as measured by transmission electron microscopy (Supporting Information, **Fig. S3**).^54,55^ UV-Vis spectra were recorded at 20 °C with an Olis-refurbished, Peltier-controlled Agilent 8453 UV-Vis Spectrophotometer in the 250-700 nm range using 1-cm path length quartz cuvettes. The UV-Vis spectrum of AuNP shows a typical intense plasmon resonance band, centered at 520 nm (Supporting Information, **Fig. S4**), in the spectrum of AuNP sample.^56^ The concentration was determined by its UV-vis peak absorption at 520 nm using a molar absorptivity of 3.94 ×10^8^ M^-1^ cm^-1^.^57^ The concentrated AuNPs were stored at 4 °C.

### Preparation of Elastin-Like Polypeptide (ELP) Constructs

DNA sequences were designed to encode elastin like sequence [VPGXG]40 (X is the guest residue) and purchased from GeneArt gene synthesis (Thermo Fisher Scientific Inc., Waltham, MA). The sequence used corresponds to ELP40, a 40-repeat ELP containing mainly VPGVG repeats (Supporting Information, **Fig. S5**). Cysteine residues were introduced either at position 4 (A4C) or 84 (A84C) to facilitate AuNP attachment. The sequences were cloned into a pET25b vector and expressed in *E. coli* BLR (DE3) cells. The expression strains were grown by inoculating cells in terrific broth (TB) medium. The purification was performed as described previously.^23,27,28^ For the cysteine variants, dithiothreitol (DTT) was added to keep the cysteine in reduced form during the purification process. In the final step, all the proteins were dialyzed into the fresh buffer containing 20 mM HEPES, 5 mM NaCl and 5 mM TCEP. Purified proteins were lyophilized for long-term storage after dialyzing the proteins in milli-Q water.

### Preparation of GB3 Null Fusion (GB3NF) Construct

The GB3NF variant was designed by mutating N8C for GB3 to form a covalent bond between the 8th residue and the AuNP surface. The plasmid was purchased from Thermo Fisher Life Technologies. This sequence contains the GB3 protein, a 6X histidine tag to aid in purification a thrombin cleavage site allowing the His tag to be removed, and a sequence of flexible Gly-Ser repeats representing a “null” fusion protein (Supporting Information, **Fig. S5**). The pET-15b vector was used to subclone the DNA for expression in *E. coli* BL21 (DE3) cells. The transformed cells were grown by inoculation in a starter culture LB media continuing 100 μg mL^-1^ ampicillin overnight at 37 °C. This culture was then added to 1L of media to bring the initial OD600 to 0.05. The larger culture was incubated at 37 °C in a 200-rpm shaker. When this culture reached an OD600 of 0.5– 0.7, expression was induced with a final concentration of 1 mM isopropyl β-D-1-thiogalactopyranoside (IPTG) and harvested after 6 h. These cells were pelleted by centrifugation for 30 min at 7000 × *g*, and then re-suspended in lysis buffer (50 mM NaCl, 20 mM NaH2PO4 pH 4.5, 5 mM EDTA, 0.5 mg mL^-1^ lysozyme, 1 mM DTT). The resuspended cells were sonicated on ice in a Thermo-Fisher sonicator 250 at 45 % power level for 3 min of total processing time (30 seconds pulse, 30 seconds rest). After lysis, solution was incubated in an 85°C hot water bath for 15 minutes to allow other cellular proteins to unfold and precipitate, with swirling every 3-4 min. Afterward, the solution was immediately transferred to an ice water bath, and streptomycin sulfate (final concentration 1 mg mL^-1^) was added to precipitate DNA. The denatured proteins remaining in the solution were removed by centrifugation at 18,000 x *g* for 45 min. GB3NF remaining in the soluble fraction was purified on an AKTA purification system connected with a HiTrap QFF anion exchange column for purification (Cytiva). The flow through with wash buffer (50 mM NaCl, 20 mM NaH2PO4 pH 4.5) was collected. The collected fractions were concentrated using 3.0 kDa molecular weight cutoff centrifugal filters.

The purified protein was dialyzed in a wash buffer containing thrombin to cleave the His-Tag at the N-terminus. Benzamidine FF beads (Cytiva) were then used to remove thrombin. The final purification was done with a HiLoad 26/600 Superdex 75 pg gel filtration column (Cytiva). The eluted proteins were pooled and dialyzed in dialysis buffer (20 mM HEPES, 5 mM NaCl, 5 mM TCEP and pH6.5). The molecular weight and purity of the GB3NF was confirmed by SDS-PAGE.

### Preparation of Temperature-Responsive Nanospheres (TRNs)

AuNP@ELP were prepared by diluting ELP in dialysis buffer to the appropriate concentration (see text for protein: AuNP ratios), and then the appropriate volume of concentrated AuNPs was added from a stock solution to bring the final volume to 1 mL. Preparation of AuNP@ELP@GB3NF was done in similar manner, where desired volumes of ELP and GB3NF were pre-mixed in dialysis buffer and then mixed with AuNPs to bring the final volume to 1 mL. After incubation for two hours, the solutions were centrifuged three times at 21,300 x *g* for 24 min to pellet AuNPs and remove residual proteins in the supernatant. The pH of the prepared solutions was confirmed to remain at 6.5 from the supernatant of the first wash with pH-indicator strips (EMD Millipore). After the third wash the pellet was resuspended in sterile 20 mM HEPES, 100 mM NaCl, and pH 6.5 to provide a suitable environment for phase separation. AuNPs were modified with HS-PEG to form AuNP@PEG5K as a control^51^.

### Quantifying Total Gold Concentration

Inductively coupled plasma mass spectrometry (ICP-MS) was used to determine the total gold content in each particle sample. TRNs were treated for 12 hr with aqua regia prior to dilution to a final volume of 10 mL with ultrapure water. A standard curve was generated consisting of at least five points, bracketing the anticipated total gold content. A commercial gold standard (Fluka #38168) was carefully diluted to generate standard curves. Samples were run on a Perkin Elmer ELAN DRC II ICP-MS system with autosampler. Total gold content (ppm, or μg mL^-1^) was determined by interpolation in the standard curve. To facilitate comparisons with other work, we have reported concentrations both as the particle concentration estimated using the extinction coefficient (nM) as well as the total concentration of gold atoms determined by ICP-MS (ppm).

### Characterization of Thermal Responsiveness

DLS was used to analyze the self-association of the TRNs at different temperatures. Quartz cuvette (Starna cells, # 16.45F-Q-3/Z8.5) containing 50 μL of the samples at 10 nM particle concentration (120 ppm total Au) concentration were incubated from 30 to 55 °C, and the hydrodynamic diameter (DH) was monitored every 0.2 °C near the phase separation temperature (T_t_). The DLS cuvette was inspected for air bubbles prior to the experiment and 50 μL of paraffin oil was added above the sample to prevent evaporation during the DLS measurements. At each temperature, the system was equilibrated approximately 3 min prior to the measurements. For reversibility experiments of AuNP@ELPA4C and AuNP@ELPA4C@GB3NF, the system was equilibrated for 15 min at 20 °C and 2-3 °C above the T_t_ for 5 cycles. The AuNP@ELPA84C and AuNP@ELPA84C@GB3NF materials were slower to return to solution, so for these samples the system was equilibrated 15 min at 20 °C and 2-3 °C above the T_t_ for only 1 cycle. Then the samples were kept at room temperature for 24 h and the size was measured.

UV-Vis absorption spectroscopy measurements were performed using a Olis-refurbished Agilent 8453 UV-Vis Spectrophotometer equipped with a Quantum Northwest integrated Pelteier temperature controller. The T_t_ were measured using a quartz cuvette with 1 cm path length. At each temperature, the system was equilibrated approximately 3 min prior to the experiments. The measurements were carried out using 500 μL of the samples (2 nM particle concentration, or 24 ppm total Au) at a rate of 0.2 °C/3 min. For reversibility experiments of AuNP@ELPA4C and AuNP@ELPA4C@GB3NF, the system was equilibrated 15 min at 20 °C and 2 −3 °C above the T_t_ for 5 cycles.

### Transmission Electron Microscopy (TEM) Measurements

Aliquots of 5 µL of 2 nM (24 ppm Au) AuNP and AuNP@ELP solution was deposited on Formvar-coated copper grids. The excess liquid was wicked away, and the remaining thin film on the grid was allowed to dry for 24 h. at room temperature or 2-3 °C above the T_t_. Prepared grids were imaged using JEOL 2100 with an accelerating voltage of 200 kV. TEM was performed at the Institute for Imaging and Analytical Technologies (I^2^AT) at Mississippi State University.

### Photothermal Conversion Efficiency Measurements

AuNP@ELP samples at different gold concentrations in a disposable cuvette was incubated 1-2 °C above the T_t_ for 15 min and then exposed to irradiation at 808 nm laser (Dragon Lasers) at 1.2, 1.4, 1.6, and 1.8 W cm^-2^. Power output was determined by the manufacturer. The sample temperatures were recorded using an infrared thermographic camera every 100 ms (Optris PI400i). To calculate the photothermal conversion efficiency (ɳ) of TRNs, 1 mL of 20 nM particles (240 ppm total Au) were incubated at their T_t_ for 15 min and then exposed to laser irradiation (808 nm, 1.8 W). When the temperature stabilized at a maximum, the laser was switched off. The parameter ɳ was calculated using the following equations:^3,33,58^

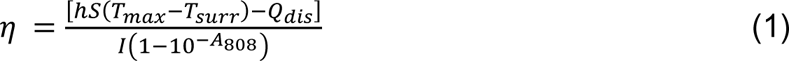

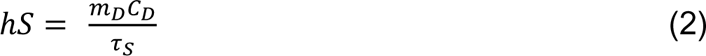

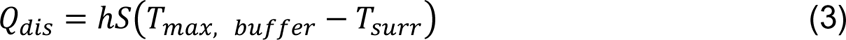

In these equations, ℎ is the heat transfer coefficient and *S* is the surface area of the container. Since these parameters are challenging to determine precisely, they are estimated from the thermal time constant (*τ_s_*), the mass of the system (*m*_*D*_, 1 gm), and the specific heat capacity of the solvent (*C*_*D*_, 4.186 J g^-1^ K^-1^). *T*_*max*_ and *T*_*max, buffer*_ are the maximum steady state temperatures with and without TRNs, respectively. *T*_surr_ is the temperature of the surrounding environment. *Q*_*dis*_ is heat dissipated from the light adsorbed by the solvent and the container, *I* is the laser power, and *A*_808_ is the UV-Vis extinction of the TRNs at 808 nm, measured at 1 cm path length (the same path length experienced by laser irradiation). To calculate the ℎ*S* via equation 2, the cooling curve was monitored after laser irradiation was removed. The plot of temperature vs. time during cooling was used to calculate the value of *τ_s_* two ways: In the first, *τ_s_* was fit using nonlinear least squares in the equation *T* = *T*_0_*e*^−*t*/*τ_s_*^ + *C*. Only the cooling decay points were used in this fit. In the second approach, *τ_s_* was determined as the average slope of the plot of ln *T* vs. time.^58^ Of the two methods, the first was found to yield more reproducible results.^59^ Obtaining *τ_s_* enabled the determination of ℎ*S*, and from this, *η* and *Q*_*dis*_ could be calculated via equations 1 and 3.

### Cell Culture and Maintenance

Human embryonic kidney (HEK-293) cell line (passage 5) was procured from the American Type Culture Collection (ATCC^®^ CRL-2302^™^) (Manassas, Virginia, USA) and was maintained in DMEM medium with 10% heat-inactivated FBS, and Antibiotic-Antimycotic (100 μg mL^-1^). Cells were maintained at 37 °C in a humidified atmosphere with 5% CO2. The culture medium was changed every 2-3 days. All experiments were conducted on cells between passages 5 to 10.

### Cytocompatibility Assay

The glow assay (CellTiter Glo) was used to examine the cytocompatibility of various gold nano formulations. Briefly, 5 × 10^3^ per well of HEK-293 cells were seeded into a 96-well plate with 200 μL of DMEM medium (10% FBS+1% Antibiotic). The 96-well plate containing seeded cells was incubated overnight at 37 °C with 5% CO2. The medium was restored with 200 μL of fresh medium containing PBS or various concentrations (10, 20 and 40 nM particle concentration, or 120, 240, or 480 ppm total Au) of AuNP, AuNP@PEG5K, AuNP@ELPA84C and AuNP@GB3NF. The treated cells were incubated for 24 h and then washed three times with PBS. After washing, wells were refreshed with 100 μL fresh medium and 100 μL CellTiter-Glo® reagent (equal to the volume of cell culture medium) and agitated for 2 min on an orbital shaker to induce cell lysis. Then, the plate was incubated at room temperature for 10 min to stabilize the luminescent signal, and luminescence was recorded using a microplate reader (Bio Tek, USA). The experiments were performed in triplicate. The results were compared with the control (no treatment) cell viability.

### In vitro Antibiofilm Activity of TRNs Irradiated by NIR Light

*S. epidermidis* strain 1301 (a gift from Dr. Keun Seo, College of Veterinary Medicine, MSU) and *E. coli* strain ATCC 25992 (a gift from Dr. Justin Thornton, Department of Biological Sciences, MSU) were employed in this study. *S. epidermidis* and *E. coli* were inoculated from a stab culture into Brain Heart Infusion (BHI) media at 37 °C and kept overnight with shaking for growth. For culturing, 100 μL of seed culture with 0.2 OD600 was added to 96-well plates. The plate was incubated statically at 37°C for 48 h, allowing biofilms to form. Following the incubation and removal of the excess cells with three washes of 100 μL of sterile PBS buffer by pipetting. A crystal violet assay and scanning electron microscopy (SEM) were used to evaluate the biofilm formation (Supporting information, **Fig. S14**). After the biofilm was obtained, the wells were treated with 100 μL of TRNs and kept 1-2 °C above the T_t_ for 15 min and then exposed to irradiation at 808 nm laser at 1.8 W cm^-2^ for 5 min.

### Determination of Bacterial Culture Densities

After NIR irradiation, the TRNs were removed from each well of a 96-well plate and 100 μL of sterile PBS was added into each well. Then each well was vigorously pipetted to disperse the biofilms. The suspensions were 10X serially diluted with sterile PBS, and 10 μL diluted samples were spread on agar plates. A total of 10 serial dilutions were performed. After incubation at 37 °C for 18 h, the colonies were counted. The number of colony forming units per mL (CFU mL^-1^) was determined using the dilution factor and number of discrete colonies observed when more than 10 colonies were present in a given dilution.

## Supporting information

All Supporting Information

## Associated Content

### Supporting Information

The supporting information is available free of charge at … Experimental characterization of AuNPs, additional temperature response curves, control experiments involving PEG5K, images of bacterial biofilms, additional experiments documenting anti-biofilm activity, cytotoxicity assays, and example calculations. (PDF)

All raw data has been uploaded to Zenodo at https://doi.org/10.5281/zenodo.8242125.

## Author Contributions

D.L.A, T.A.W., and N.C.F designed experiments. D.L.A., C.S.K, M.A.H, V.B.T, T.A.W., and N.C.F performed experiments. D.L.A. and N.C.F. wrote the initial draft. D.L.A., C.S.K, M.A.H, V.B.T, T.A.W., and N.C.F revised the manuscript. All authors agreed to the final manuscript.

### Funding Sources

This work was supported by the National Institutes of Health under award number R01AI139479 (NCF) and the National Science Foundation under award 1818090 (NCF), as well as the American Cancer Society under award RSG-21-114-01-MM (TAW).

### Notes

The authors declare no competing financial interest.

## Acknowledgements

We thank G. Lee Bidwell III for helpful suggestions in the initial planning of this project and Jack Correia for critical reading of the manuscript.

## Abbreviations

AuNP: gold nanoparticle
Ang II: angiotensin II
AuNR: gold nanorod
AuNS: gold nanosphere
DH: hydrodynamic diameter
DLS: dynamic light scattering
ELP: elastin-like polypeptide
GB3NF: GB3 protein “null” fusion
LSPR: localized surface plasmon resonance
PEG: polyethylene glycol
PTT: photothermal therapy
TRN: temperature-responsive nanosphere
T_t_: transition temperature

