## Supplementary material for "Near-Infrared Photothermal Ablation of Biofilms using Protein-Functionalized Gold Nanospheres with a Tunable Temperature Response": All Supporting Information

**Supporting Methods: Photothermal Conversion Efficiency (PCE) Calculations**

Values are shown for the AuNP@ELPA84C (1:200) construct (see **Fig. 4E**) and AuNP@PEG. The equations used are:[1–3]

$$\eta = \frac{[hS(T_{max} - T_{surr}) - Q_{dis}]}{I(1 - 10^{-A_{808}})}$$

$$Q_{dis} = hS\,(T_{max,\ buffer} - T_{sur})$$

$$\frac{m_D C_D}{hS} = \tau_s$$

From the cooling curve, the value of $\tau_S$ was determined as 370 s (see Fig. 4E):

$$\frac{m_D C_D}{hS} = 370\ s$$

In this equation $m_D$ was 1.0 g, and the specific heat capacity ($C_D$) was 4.186 $J\ g^{-1}K^{-1}$. The value of $Q_{dis}$ was calculated using an experiment without AuNPs (see Fig. 4E inset). $A_{808}$ was 0.1493.

**PCE of AuNP@ELPA84C above the transition temperature.**

$$\eta = \frac{\left[\frac{(1.0\ g)(4.186\ J\ gm^{-1}K^{-1})}{370\ s}(37.8 - 6.0K)\right]}{(1.8)(1 - 10^{-0.1493})}$$

$$\eta = \frac{0.1998}{(1 - 0.7090)}$$

$$\eta = \frac{0.1998}{(0.2909)}$$

=0.6868

=68.7%

**PCE of AuNP@ELPA84C below the transition temperature.**

$$\eta = \frac{\left[\frac{(1.0\ g)(4.186\ J\ gm^{-1}K^{-1})}{370\ s}(31.0 - 6.0K)\right]}{(1.8)(1 - 10^{-0.1493})}$$

$$\eta = \frac{0.1571}{(1 - 0.7090)}$$

$$\eta = \frac{0.1571}{(0.2909)}$$

=0.5401

=54.01%

**PCE of AuNP@PEG5K.**

$$\eta = \frac{\left[\frac{(1.0\ g)(4.186\ J\ gm^{-1}K^{-1})}{405\ s}(23.69 - 6.0K)\right]}{(1.8)(1 - 10^{-0.1493})}$$

$$\eta = \frac{0.1014}{(1 - 0.7090)}$$

$$\eta = \frac{0.1014}{(0.2909)}$$

=0.348

=34.8%

**Supporting Figures**

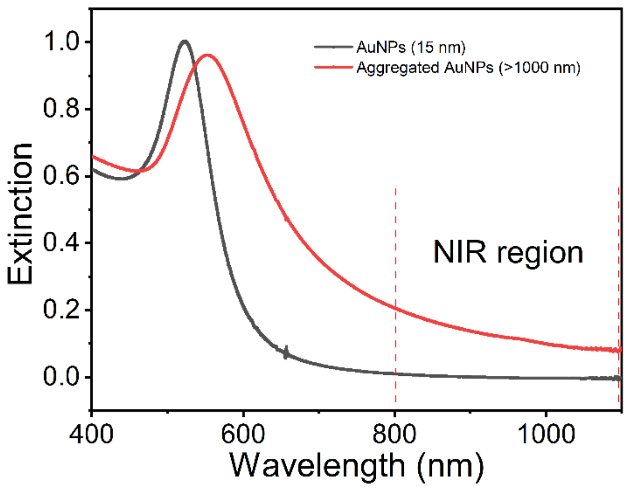

**Figure S1. Effect of NIR absorption on the size of the AuNPs.**

Data is shown for synthesized 15 nm AuNPs (black curve) and thermally agglomerated AuNP@ELPA84C (1:200) at 39 ℃ (red curve)

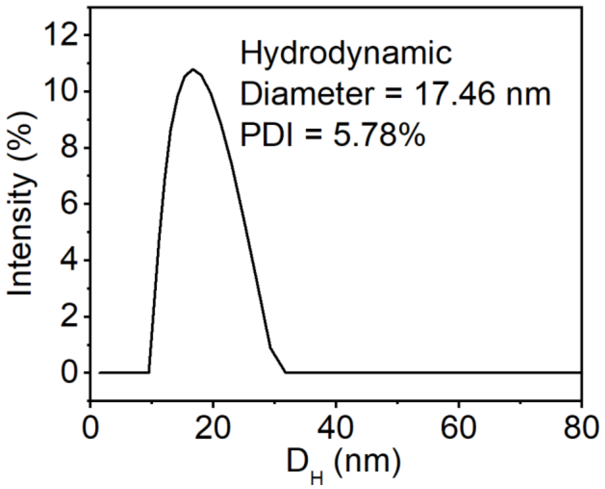

**Figure S2. Particle size distribution of synthesized AuNPs.**

Data is shown for particle size distribution of synthesized 15 nm AuNPs measured using Anton Paar Dynamic Light Scattering (DLS) system at room temperature.

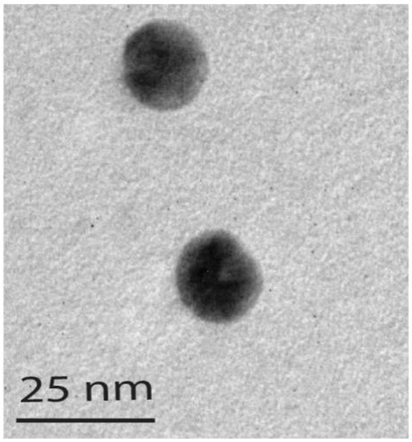

**Figure S3. TEM image of AuNPs.**

Data is shown for synthesized 15 nm AuNPs.

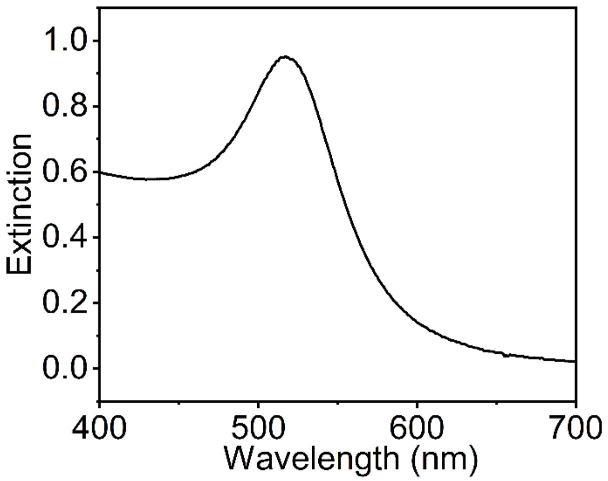

**Figure S4. Extinction spectrum of AuNPs.**

Data is shown for the extinction of synthesized 15-nm AuNPs as a function of wavelength.

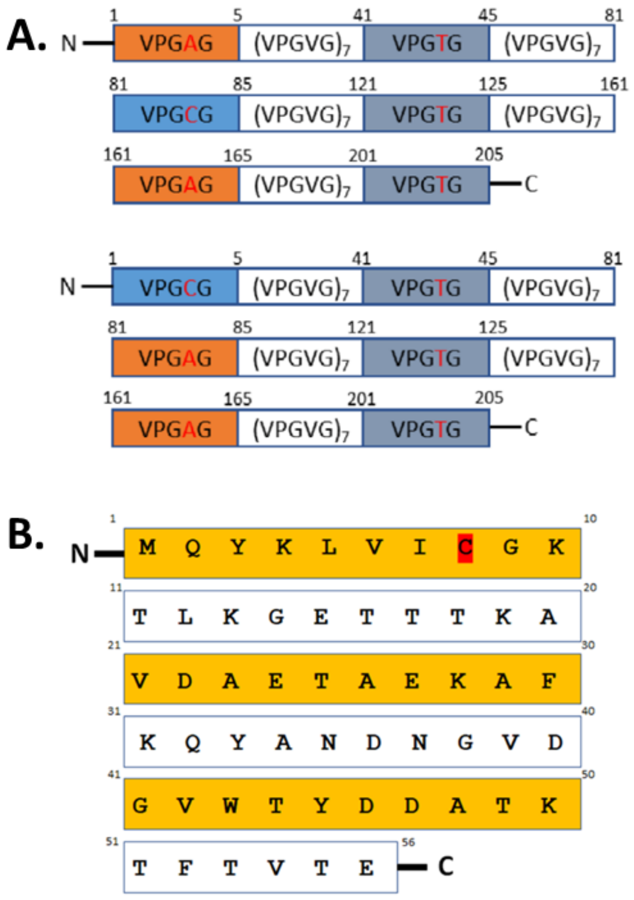

**Figure S5. Amino acid sequence of ELP and GB3NF.**

Schematic representation of ELPA84C (top) and ELPA4C (bottom) amino acid sequences (A). Sequence of GB3 null fusion (GB3NF, B).

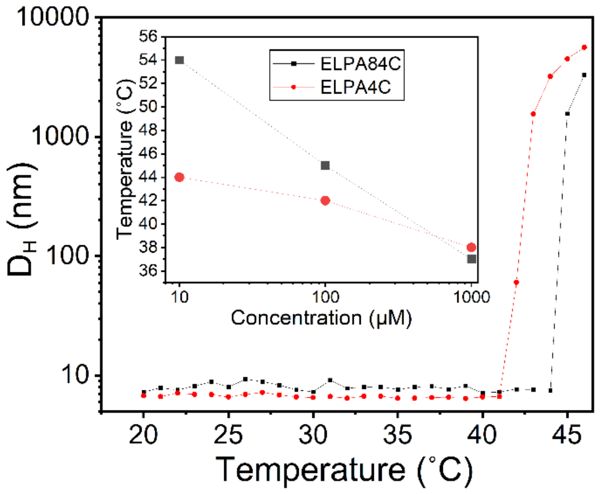

**Figure S6. Temperature-induced liquid-liquid phase separation (LLPS) of ELPs**

Data is shown for inverse thermal transition (the agglomeration of ELP at high temperatures) of ELPA4C in red and EPA84C in black. $T_t$ was concentration-dependent (inset).

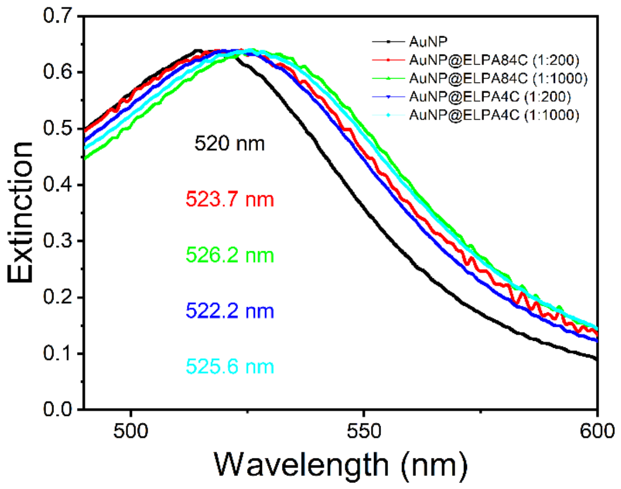

**Figure S7. UV-Vis profiles of TRNs.**

Data is shown for extinction of bare AuNP (black), AuNP@ELPA84C (1:200) (red), AuNP@ELPA84C (1:1000) (green), AuNP@ELPA4C (1:200) (blue) and AuNP@ELPA4C (1:1000) (cyan). All spectra are normalized.

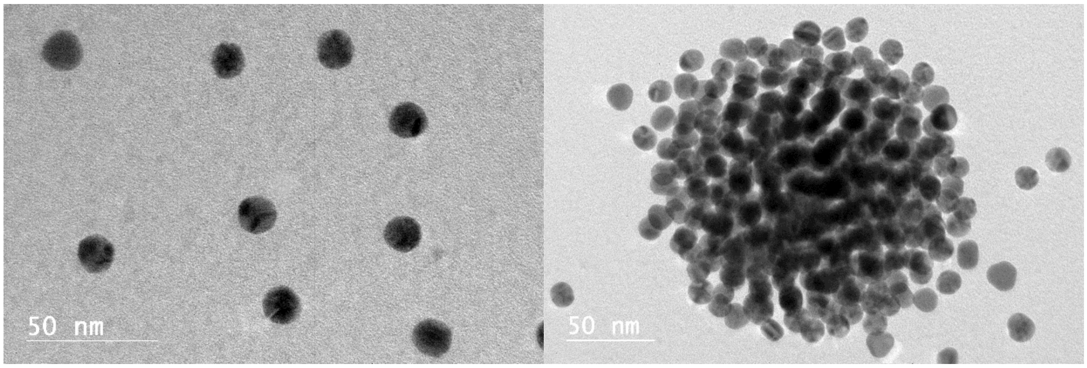

**Figure S8. TEM images representing temperature-responsiveness of AuNP@ELP.**

Data is shown for AuNP@ELPA4C below the transition temperature (left) above the transition temperature (right).

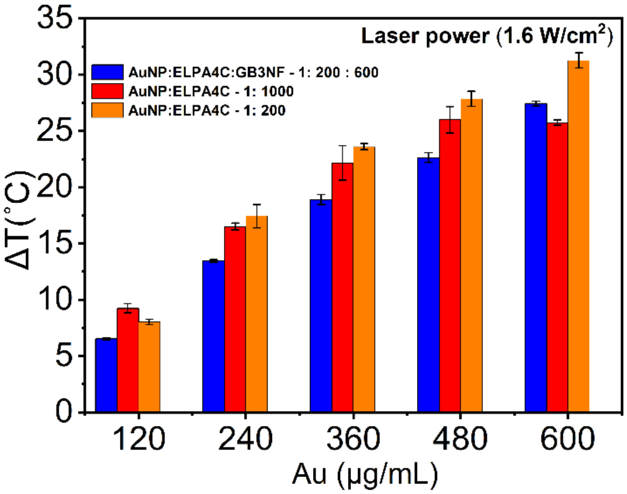

**Figure S9. Gold concentration dependence on photothermal effect of TRNs.**

The effect of gold concentration on ELPA4C based TRNs when exposed to laser irradiation 808 nm (1.6 W, 5 min), after incubation for 15 min at their transition temperatures ($T_t$).

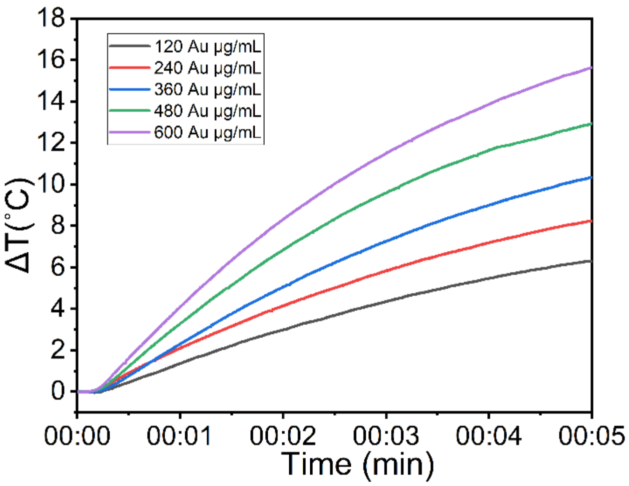

**Figure S10. Gold concentration dependence on photothermal effect of AuNP@PEG5K.**

Heating curves of AuNP@PEG5K at different concentrations of gold under 808 nm laser irradiation (1.6 W, 5 min), after incubation for 15 min at 35 °C. ΔT denotes the temperature evolution.

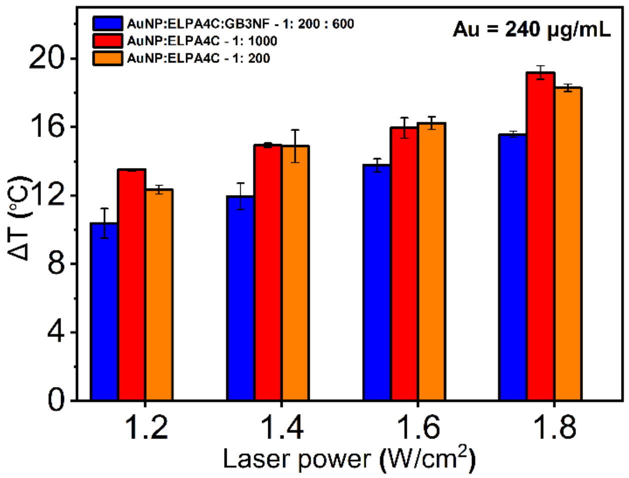

**Figure S11. Laser power dependence on photothermal effect of TRNs.**

The effect of laser power on ELPA4C based TRNs when exposed to laser irradiation 808 nm (5 min), after incubation for 15 min at their transition temperatures ($T_t$).

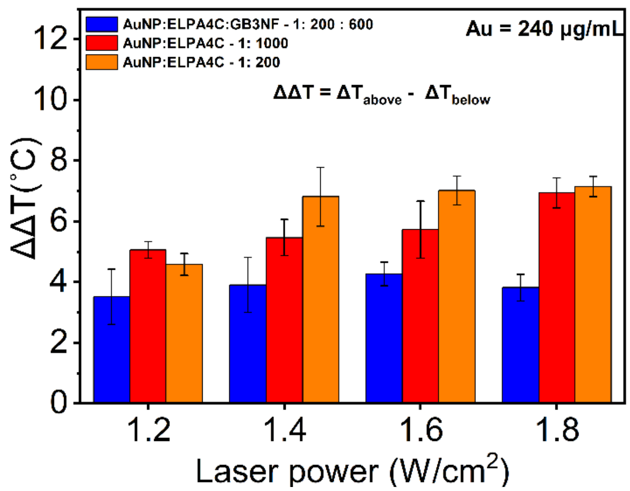

**Figure S12. Temperature dependence on photothermal effect of TRNs.**

The effect of agglomerated vs. non-agglomerated state of ELPA4C based TRNs when exposed to laser irradiation 808 nm (5 min), after incubation for 15 min at their transition temperatures ($T_t$).

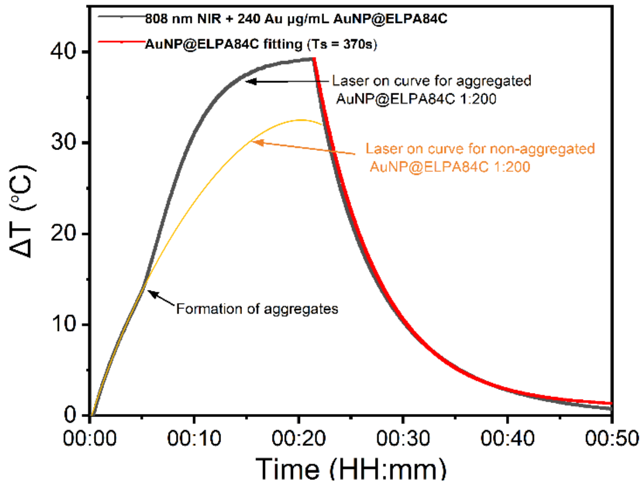

**Figure S13. Photothermal effect of AuNP@ELPA84C.**

Laser on and off curves for AuNP@ELPA84C (1:200), when 20 nM (Au 240 μg/mL) aqueous solution was irradiated with a 1.8 W/cm$^2$, 808 nm laser at 22 ℃. The change in logarithmic growth curve around 8 min denotes the formation of nanoparticle agglomerates.

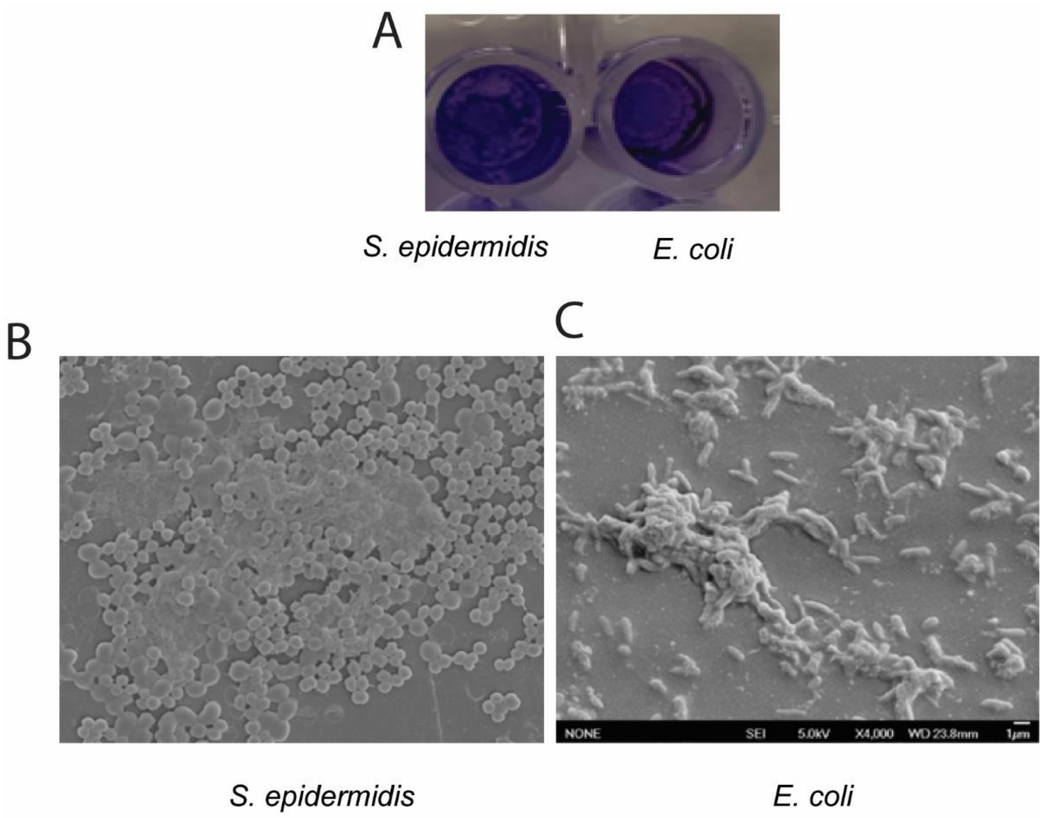

**Figure S14. Images *S. epidermidis* and *E.coli* biofilm formation on polystyrene surfaces.**

(A) Crystal violet staining of *S. epidermidis* (Strain 1301, left) and *E.coli* (ATCC 25992, right) biofilm on 96-well plate. SEM images of (B) *S. epidermidis* (Strain 1301) (C) *E.coli* (ATCC 25992) on polystyrene surfaces.

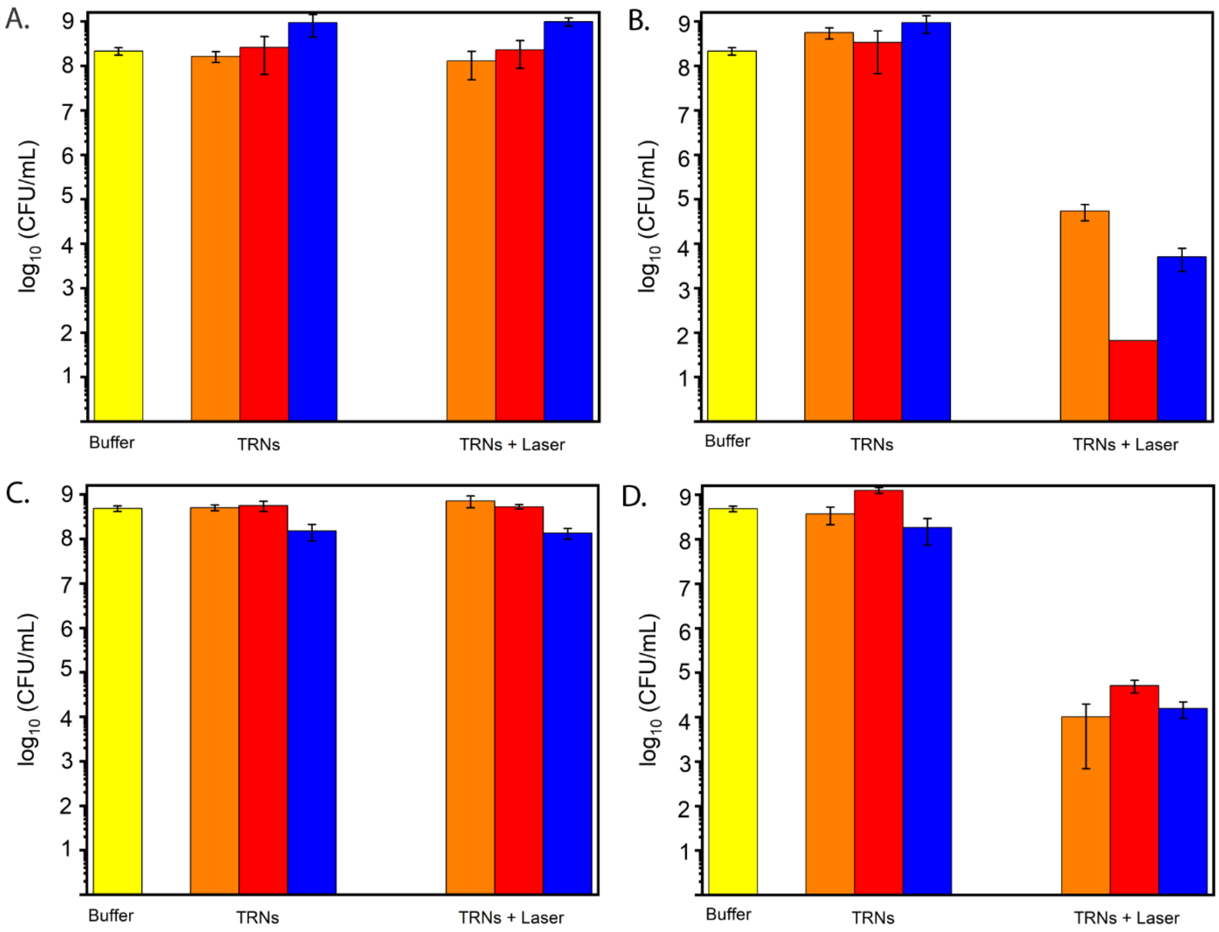

**Figure S15. In vitro antibacterial activities of TRNs against antibiotic-sensitive bacteria.**

Statistical analysis of bacterial cell viability by $\log_{10}$ (CFU mL $^{-1}$). (A). *E. coli* at temperatures below $T_t$. (B). *E. coli* at temperatures above $T_t$. (C). *S. epidermidis* at temperatures below $T_t$. (D). *S. epidermidis* at temperatures above $T_t$. . [Yellow: Buffer (20 mM HEPES, 100 mM NaCl, pH 6.5), Orange: AuNP@ELPA4C (1:200), Blue: AuNP@ELPA4C@GB3NF (1:200:600), and Red: AuNP@ELPA4C (1:1000)] Error bars represent the standard error of the mean for at least three independently prepared TRN samples.

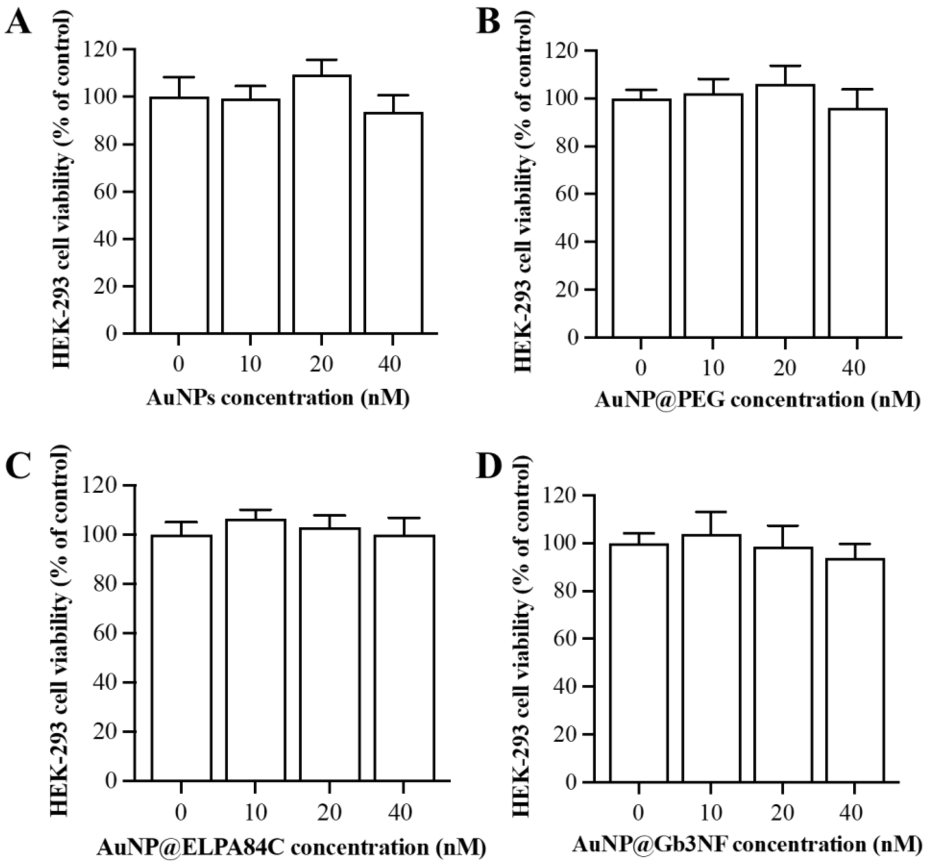

**Figure S16. Effects of different AuNP formulations on the viability of HEK-293 cells.**

Effects of (A) AuNPs (B) AuNP@PEG5K, (C) AuNP@ELPA84C and (D) AuNP@GB3NF on cell survival rate of the HEK-293 cells. Data are expressed as the mean ± standard deviation. Treatment group changes were represented with respect to percentage of control.

### Supporting Information References

(1) Park, S.; Lee, W. J.; Park, S.; Choi, D.; Kim, S.; Park, N. Reversibly PH-Responsive Gold Nanoparticles and Their Applications for Photothermal Cancer Therapy. *Scientific Reports* **2019**, *9* (1). https://doi.org/10.1038/s41598-019-56754-8.

(2) Yuan, Z.; Lin, C.; He, Y.; Tao, B.; Chen, M.; Zhang, J.; Liu, P.; Cai, K. Near-Infrared Light-Triggered Nitric-Oxide-Enhanced Photodynamic Therapy and Low-Temperature Photothermal Therapy for Biofilm Elimination. *ACS Nano* **2020**, *14* (3), 3546–3562. https://doi.org/10.1021/acsnano.9b09871.

(3) Lin, M.; Guo, C.; Li, J.; Zhou, D.; Liu, K.; Zhang, X.; Xu, T.; Zhang, H.; Wang, L.; Yang, B. Polypyrrole-Coated Chainlike Gold Nanoparticle Architectures with the 808 Nm Photothermal Transduction Efficiency up to 70%. *ACS Applied Materials & Interfaces* **2014**, *6* (8), 5860–5868. https://doi.org/10.1021/am500715f.
